# MosaiCatcher v2: a single-cell structural variations detection and analysis reference framework based on Strand-seq

**DOI:** 10.1101/2023.07.13.548805

**Authors:** Thomas Weber, Marco Raffaele Cosenza, Jan Korbel

## Abstract

Single-cell DNA template strand sequencing (Strand-seq) allows a range of various genomic analysis including chromosome length haplotype phasing and structural variation (SV) calling in individual cells. Here, we present MosaiCatcher v2, a standardised workflow and reference framework for single-cell SV detection using Strand-seq. This framework introduces a range of functionalities, including: an automated upstream Quality Control (QC) and assembly sub-workflow that relies on multiple genome assemblies and incorporates a multistep normalisation module, integration of the scNOVA SV functional characterization and of the ArbiGent SV genotyping modules, platform portability, as well as a user-friendly and shareable web report. These new features of MosaiCatcher v2 enables reproducible computational processing of Strand-seq data, which are increasingly used in human genetics and single cell genomics, towards production environments.

**Availability and Implementation:** Mosaicatcher v2 is a standardised workflow, implemented using the Snakemake workflow management system. The pipeline is available on GitHub: https://github.com/friendsofstrandseq/mosaicatcher-pipeline/ and on the snakemake-workflow-catalog:

https://snakemake.github.io/snakemake-workflow-catalog/?usage=friendsofstrandseq/mosaicatcher-pipeline.

**Supplementary information:** Supplementary data are available at Bioinformatics online.

## 1. Introduction

Strand-seq is an amplification-free single-cell short-read sequencing technique that generates strand-specific libraries by targeting and sequencing specifically template strand during DNA replication (Falconer et al. 2012). Strand-seq methodology enables diverse applications not readily accessible to other technologies: characterization of a wide variety of SV classes in single cells including balanced inversions as well as complex genomic variation such as chromothripsis events (Sanders et al. 2016; Porubsky et al. 2022), locating sister chromatid exchange events (Claussin et al. 2017), single cell multi-omic analyses linking genome and molecular phenotype in the same cell (Jeong et al. 2022), generation of long-range phase information to assist genome assembly (Porubsky et al. 2020; 2022; Ebert et al. 2021; Jarvis et al. 2022), contiguity validation of long-range assemblies (Nurk et al. 2022), and characterisation of somatic SV events at the single-cell level, and thus in subclonal cell populations, to foster studies in cancer evolution (Sanders et al. 2020; Jeong et al. 2022)

Regarding this latter application, a computational framework, named MosaiCatcher was developed alongside the wet lab methodology in order to process Strand-seq data, based on a tri-channel processing implementation including three layers of information: depth of coverage, strand directionality and haplotype phase (Sanders et al. 2020). Here, we present MosaiCatcher v2, addressing both prior technical limitations and providing a range of new functionalities including: an upstream QC and assembly sub-pipeline that can rely on the most recent human genome assemblies (hg19, hg38, T2T-CHM13), as well as on mouse genome assembly (mm10) and a new multistep normalisation module. Additionally, MosaiCatcher v2 incorporates a structural variation (SV) functional analysis module, which uses nucleosome occupancy data measured directly from Strand-seq libraries (scNOVA) as well as a SV genotyper (ArbiGent). This updated version also offers increased stability through data-conditional dependent execution, and a sharable user-friendly HTML web report including all the relevant outputs, statistics and plots computed during the workflow execution. MosaiCatcher v2 is a standardised, stable and portable workflow designed to serve as a core pipeline for researchers working with Strand-seq data. Our primary goal here is to streamline and enhance the efficiency of processing Strand-seq data for the scientific community. By incorporating the latest advancements, MosaiCatcher v2 not only offers an easy-to-use pipeline but also ensures compatibility with high-scale production environments, thus enabling researchers to tackle large-scale studies and accommodate the growing volume of Strand-seq data using a comprehensive, integrated system.

## 2. Features

Mosaicatcher v2, as its prior release, relies on snakemake (Mölder et al. 2021), a workflow management system widely adopted in the bioinformatics community. Snakemake allows portability and scalability to almost all existing cluster and cloud platforms based on conda environments and containers. With the aim of standardising Strand-seq data analysis and promoting the dissemination of this technology, MosaiCatcher (schematically represented in Figure 1) we have improved stability and integrated new functionalities (listed below and detailed in the Supplementary Material):

**Figure 1.**
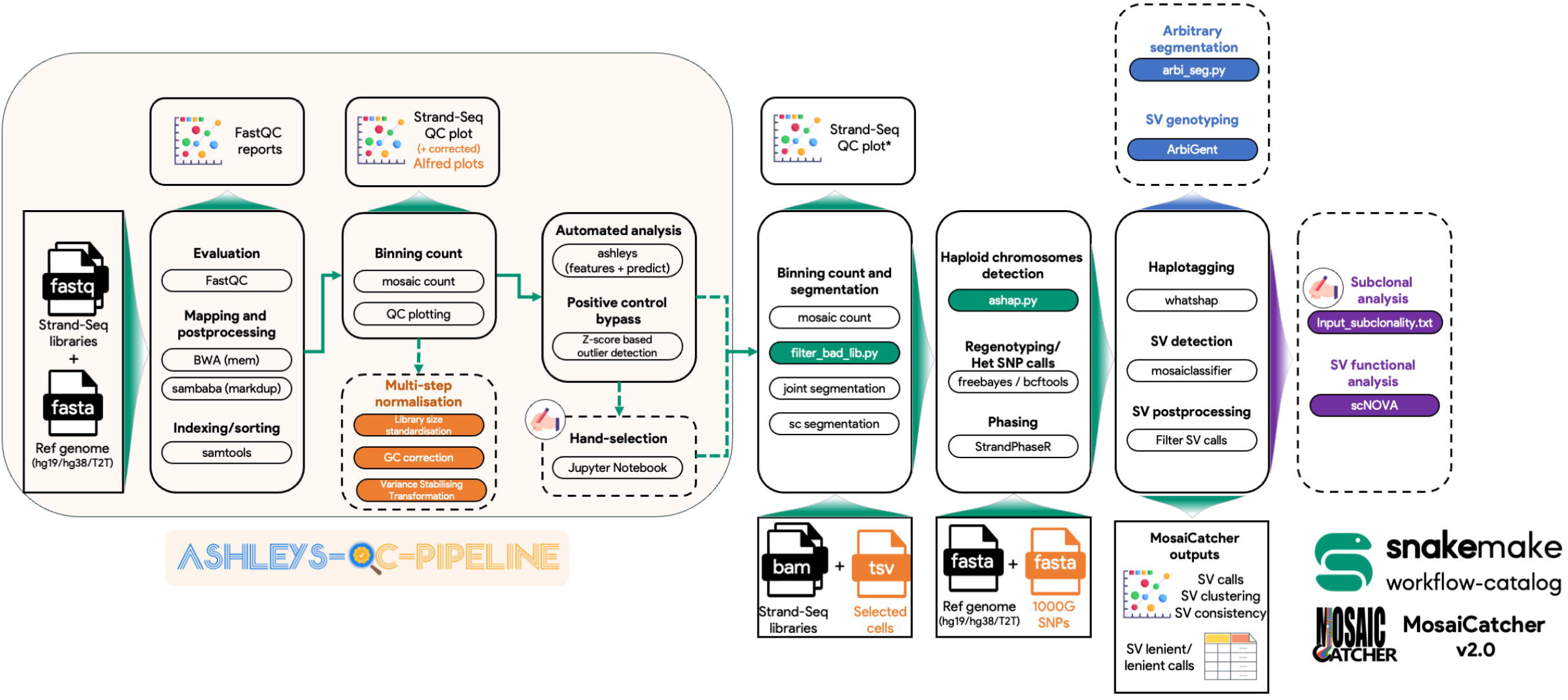
MosaiCatcher v2 schematic representation. On the left part in dimmed orange is represented ashleys-qc-pipeline, a switchable preprocessing optional module that allows to perform standard steps of mapping, sorting and indexing FASTQ libraries, producing Quality Control plots and reports as well as identifying high-quality libraries. On the right uncolored part, the MosaiCatcher core part of the pipeline is still usable as a standalone by providing Stand-Seq aligned BAM files. Green boxes correspond to data-conditional dependent execution steps (snakemake checkpoints) that allow more flexibility and reduce issues when executing the workflow. Orange box corresponds to the multi-step normalisation module. Blue box corresponds to ArbiGent mode of execution that allows SV genotyping from arbitrary segmentation. Violet box corresponds to scNOVA SV function analysis mode of execution. Dashed boxes correspond to optional modules. * : Strand-seq QC plot are only produced here if ashleys-qc pipeline is not enabled

### Strand-seq end-to-end processing and analysis

the previous version of MosaiCatcher relied on cumbersome manual preprocessing and filtering of high-quality libraries. To address this issue, the ashleys-qc (Gros et al. 2021) machine-learning-based tool was developed to automatically select high-quality libraries. In this new MosaiCatcher version, ashleys-qc is seamlessly integrated, enabling end-to-end computational processing from the raw sequencing FASTQ data to the downstream analysis including single-cell SV calling and visualisation analysis, thus promoting reproducibility and reducing bias in the analysis. This is achieved using data-dependent conditional execution steps (see below) that lead to the selection and the processing of high-quality labelled libraries through ashleys-qc. However, if needed, manual filtering by a domain-expert is still possible through a switchable option that gives to the user the possibility to identify and flag the high-quality libraries through a jupyter notebook. The ashleys-qc-pipeline was cleaned and formatted to be referenced as a standardised usage pipeline on the snakemake-workflow-catalog and usable as a module, loadable in MosaiCatcher v2.

### Data-dependent conditional execution steps

to increase workflow stability and reproducibility, we implemented several data-dependent conditional execution steps, as snakemake checkpoints. These dynamic steps help to solve the agnostic view of the snakemake workflow engine that initially needs to generate a complete Directed Acyclic Graph (DAG) of execution including all inputs (here libraries) provided by the user, regardless of their composition. Thus, the first checkpoint defined allows MosaiCatcher v2 to automatically filter out libraries flagged as low-quality using both: ashleys-qc predictions and quality control measures based on reads mapping quality and coverage biases across chromosomes. The second checkpoint defined computes the average chromosomal ploidy status at the sample level (using all libraries processed) to perform haplotype phasing of heterozygous Single Nucleotide Polymorphisms (hetSNPs) exclusively on chromosomes that present at least a diploid status.

### Multiple Reference genomes

One limitation of the previous version of MosaiCatcher was to rely exclusively on the GRCh38/hg38 assembly. In MosaiCatcher v2, it is now possible to use different human assemblies (GRCh37/hg19, GRCh38/hg38, T2T-CHM13) through an automatic downloading and indexing step of the selected reference fasta file. “Blacklists” of regions corresponding to repeated regions and mapping abnormalities (mostly telomeric and pericentromeric regions) were constructed for each assembly as described previously (Sanders et al. 2020). The possibility of using different assemblies opens up Strand-seq for analysis in other species and, as well as leveraging updated human genomic assemblies for improved SV calling and characterization.

### Read count multi-step normalisation

Strand-seq libraries, similarly to other short-read based sequencing techniques, can be affected by various types of read coverage biases, including GC and mean-variance bias. Using a multiple step approach, we process the read data with a sequence of statistical methods: library size normalisation, GC correction and variance stabilising transformation (VST) (detailed in Supplementary Material). This results in a more even and unbiased read count coverage, enabling the user to have access to considerably more readable Strand-Seq karyotype plots with improved characterisation of SVs and copy number calling even in atypical or noisy libraries.

### Standardisation, Portability and Reproducibility

to promote standardisation, we implemented Continuous Integration/Continuous development (CI/CD) control tests based on GitHub actions (Table S1). These CI/CD tests check the different functionalities implemented into newer versions of MosaiCatcher as well as code formatting and linting. Additionally, to improve standardisation, we adhered to the snakemake-workflow-catalog guidelines to comply to the “standardised usage” category of the catalog. Finally, we harmonised pipeline execution with *ad-hoc* conda environments required by the different components of the pipeline, packed into a Docker container, versioned at each new pipeline release. Taken together, MosaiCatcher v2 developed into a mature, portable and reproducible workflow.

### SV genotyping with ArbiGent

AbriGent leverages strand-seq advantage in haplotype phasing for SV validation and genotyping, as well as for surveying large populations and measuring variant frequencies. This new release of Mosaicatcher features full integration of ArbiGent (Porubsky et al. 2022) to enable genotyping of inversions as well as all other types of SVs–containing at least 500 bp of uniquely mappable sequence. Among its key functionalities, ArbiGent supports (1) fine-scaled read depth normalisation to improve accuracy in complex genomic regions, (2) methods for phase harmonisation between different datasets and (3) a number of population-based QC metrics that can help identify spurious SV calls.

### SV functional analysis with scNOVA

A unique feature of Strand-Seq data is that they can additionally be interpreted as an epigenetic readout, a feature that is leveraged by scNOVA (single-cell nucleosome occupancy and genetic variation analysis) (Jeong et al. 2022), to move from the discovery of SVs in single cells to characterisation of their functional consequences in the same cell. In this updated version of the MosaiCatcher pipeline, scNOVA was seamlessly integrated and can be launched using a single unified command.

### Interactive reports and downstream analysis

to improve reproducibility and transparency, we designed HTML static web reports that can be shared easily and joined to publications, by taking advantage of snakemake functionalities. MosaiCatcher v2 interactive reports contain all plots and information (Table S2) required to analyse a Strand-seq experiment, from the primary analysis and quality control (FASTQC, Strand-seq QC count plots, GC analysis, cells excluded from the experiment) to the downstream analysis (single-cell SV calls plots based on different stringency criteria). Additionally to the interactive web reports, a UCSC-genome browser custom file is automatically produced at the end of the workflow, which includes for each individual cell level Watson and Crick read counts, as well as SV calls. As presented in Figure S6, this allows the user a more fine-grain analysis, by investigating interactively breakpoints, orientation, or potentially associated genes and regulatory elements around the identified SVs, using the different UCSC ressources.

## 3. Application

To showcase new functionalities of MosaiCatcher v2, we reprocessed - in a unique run - four cell line samples (RPE1-WT, RPE-BM510, RPE-C7 and NA20509 Lymphoblastoid Cell Line (LCL)) previously used in the original MosaiCatcher publication (Sanders et al. 2020), (Table S3). Based on the full set of raw FASTQ libraries, we performed mapping against T2T-CHM13 reference assembly and automatic selection through ashleys-qc. Complete HTML report including quality control, multi-step normalisation related visualisations and downstream analysis outputs, is available as a Supplementary data and provides an example of this more user-friendly and easy to share report that allows SV characterisation at the single-cell level.

In a second run, using the same samples listed above, we demonstrate SV genotyping of ten copy-number imbalanced SV, previously validated through Whole Genome Sequencing (WGS) and emphasising the utility of Strand-seq single cell genotyping beyond inversions (see Supplementary Material and Table S4). All the events were accurately detected and no false positives nor false negatives were detected.

## 4. Conclusion

In conclusion, MosaiCatcher V2 is a stable, reproducible and highly user-friendly computational framework with key new functionalities that allows Strand-seq data end-to-end processing, leveraging snakemake features. This update stands to be the reference Strand-seq workflow, promoting interoperability and reusability of data within the Strand-seq scientific community. Moreover, standardisation and improvements in stability greatly increases efficiency, making this pipeline well suited for high-scale processing, matching the increasing amount of Strand-seq data produced in human genetics and cancer genomics, as well as SV characterization in population studies.

## Acknowledgements

We thank E. Benito Garagorri, H. Jeong and K. Grimes for discussions. We thank Wolfram Höps for his help during ArbiGent integration into the pipeline. We thank P. Ebert, T. Marshall and T. Christensen for their developments on the ploidy assignment. We thank N. Habermann for project support. J.O.K. acknowledges funding from European Research Council Starting (grant no. 336045) and Consolidator (grant no. 773026) grants and the National Institutes of Health (grant no. 3U41HG007497-04S1).

## Funding

The authors acknowledge funding from the European Molecular Biology Laboratory (EMBL). J.O.K. received support from the German Cancer Research Center (DKFZ) and the European Research Council (ERC Consolidator grant 773026).

## Data availability statement

Example data Zenodo Part1: https://zenodo.org/record/7696695

Example data Zenodo Part2: https://zenodo.org/record/7697329

MosaiCatcher v2 publication data interactive report: https://zenodo.org/record/8005968

## Notes

https://github.com/friendsofstrandseq/mosaicatcher-pipeline

### Competing Interest Statement

The authors have declared no competing interest.

## References

Anders, Simon, and Wolfgang Huber. 2010. “Differential Expression Analysis for Sequence Count Data.” Genome Biology 11 (10): R106. https://doi.org/10.1186/gb-2010-11-10-r106.

Anscombe, F. J. 1948. “THE TRANSFORMATION OF POISSON, BINOMIAL AND NEGATIVE-BINOMIAL DATA.” Biometrika 35 (3–4): 246–54. https://doi.org/10.1093/biomet/35.3-4.246.

Auton, Adam, Gonçalo R. Abecasis, David M. Altshuler, Richard M. Durbin, Gonçalo R. Abecasis, David R. Bentley, Aravinda Chakravarti, et al. 2015. “A Global Reference for Human Genetic Variation.” Nature 526 (7571): 68–74. https://doi.org/10.1038/nature15393.

Claussin, Clémence, David Porubský, Diana CJ Spierings, Nancy Halsema, Stefan Rentas, Victor Guryev, Peter M Lansdorp, and Michael Chang. 2017. “Genome-Wide Mapping of Sister Chromatid Exchange Events in Single Yeast Cells Using Strand-Seq.” edited by Lorraine Symington. ELife 6 (December): e30560. https://doi.org/10.7554/eLife.30560.

Ebert, Peter, Peter A. Audano, Qihui Zhu, Bernardo Rodriguez-Martin, David Porubsky, Marc Jan Bonder, Arvis Sulovari, et al. 2021. “Haplotype-Resolved Diverse Human Genomes and Integrated Analysis of Structural Variation.” Science 372 (6537): eabf7117. https://doi.org/10.1126/science.abf7117.

Falconer, Ester, Mark Hills, Ulrike Naumann, Steven S. S. Poon, Elizabeth A. Chavez, Ashley D. Sanders, Yongjun Zhao, Martin Hirst, and Peter M. Lansdorp. 2012. “DNA Template Strand Sequencing of Single-Cells Maps Genomic Rearrangements at High Resolution.” Nature Methods 9 (11): 1107–12. https://doi.org/10.1038/nmeth.2206.

Gros, Christina, Ashley D Sanders, Jan O Korbel, Tobias Marschall, and Peter Ebert. 2021. “ASHLEYS: Automated Quality Control for Single-Cell Strand-Seq Data.” Bioinformatics 37 (19): 3356–57. https://doi.org/10.1093/bioinformatics/btab221.

Harrison, Paul Francis. 2017. “Varistran: Anscombe’s Variance Stabilizing Transformation for RNA-Seq Gene Expression Data.” Journal of Open Source Software 2 (16): 257. https://doi.org/10.21105/joss.00257.

Jarvis, Erich D., Giulio Formenti, Arang Rhie, Andrea Guarracino, Chentao Yang, Jonathan Wood, Alan Tracey, et al. 2022. “Semi-Automated Assembly of High-Quality Diploid Human Reference Genomes.” Nature 611 (7936): 519–31. https://doi.org/10.1038/s41586-022-05325-5.

Jeong, Hyobin, Karen Grimes, Kerstin K. Rauwolf, Peter-Martin Bruch, Tobias Rausch, Patrick Hasenfeld, Eva Benito, et al. 2022. “Functional Analysis of Structural Variants in Single Cells Using Strand-Seq.” Nature Biotechnology, November, 1–13. https://doi.org/10.1038/s41587-022-01551-4.

Mölder, Felix, Kim Philipp Jablonski, Brice Letcher, Michael B. Hall, Christopher H. Tomkins-Tinch, Vanessa Sochat, Jan Forster, et al. 2021. “Sustainable Data Analysis with Snakemake.” F1000Research. https://doi.org/10.12688/f1000research.29032.2.

Nurk, Sergey, Sergey Koren, Arang Rhie, Mikko Rautiainen, Andrey V. Bzikadze, Alla Mikheenko, Mitchell R. Vollger, et al. 2022. “The Complete Sequence of a Human Genome.” Science 376 (6588): 44–53. https://doi.org/10.1126/science.abj6987.

Porubsky, David, Peter Ebert, Peter A. Audano, Mitchell R. Vollger, William T. Harvey, Pierre Marijon, Jana Ebler, et al. 2020. “Fully Phased Human Genome Assembly without Parental Data Using Single-Cell Strand Sequencing and Long Reads.” Nature Biotechnology, December, 1–7. https://doi.org/10.1038/s41587-020-0719-5.

Porubsky, David, Wolfram Höps, Hufsah Ashraf, PingHsun Hsieh, Bernardo Rodriguez-Martin, Feyza Yilmaz, Jana Ebler, et al. 2022. “Recurrent Inversion Polymorphisms in Humans Associate with Genetic Instability and Genomic Disorders.” Cell 0 (0). https://doi.org/10.1016/j.cell.2022.04.017.

Sanders, Ashley D., Mark Hills, David Porubský, Victor Guryev, Ester Falconer, and Peter M. Lansdorp. 2016. “Characterizing Polymorphic Inversions in Human Genomes by Single-Cell Sequencing.” Genome Research 26 (11): 1575–87. https://doi.org/10.1101/gr.201160.115.

Sanders, Ashley D., Sascha Meiers, Maryam Ghareghani, David Porubsky, Hyobin Jeong, M. Alexandra C. C. van Vliet, Tobias Rausch, et al. 2020. “Single-Cell Analysis of Structural Variations and Complex Rearrangements with Tri-Channel Processing.” Nature Biotechnology 38 (3): 343–54. https://doi.org/10.1038/s41587-019-0366-x.

