## Supplementary notes for "MosaiCatcher v2: a single-cell structural variations detection and analysis reference framework based on Strand-seq"

#### 1. Software & Workflow management

MosaiCatcher v2 computational framework, as its first version (Sanders et al. 2020), is implemented as a Snakemake workflow (v7.5.0 & v7.19.1 used during development) to facilitate reproducibility, portability and scalability (URL repository below). In this second major release, the workflow was cleaned and optimised, in order to be both compliant to the snakemake-workflow-catalog standardised usage but also to leverage most of snakemake features and add new core features detailed below. Snakemake-workflow-catalog standardisation allowed better support for further development of the pipeline but also compatibility with new incoming versions of snakemake. To do so, we implemented a list of Continuous Integration and Continuous Development (CI/CD) tests using GitHub actions (see 5. Continuous Integration/Continuous Development & GitHub actions).

Mosaicatcher git repository: https://github.com/friendsofstrandseq/mosaicatcher-pipeline

#### 2. Input and external data

##### 2.1. Input data

The pipeline can be used using two different input stages as illustrated on Figure 1. As it was in the previous version, the input can be a set of single-cell (Strand-seq) BAM files (sorted, deduplicated and indexed) from the same sample. Additionally, to that, the user can now directly give to the pipeline the raw Strand-seq FASTQ files coming from the same sample. In this case, *"ashleys_pipeline"* configuration parameter needs to be enabled, leading to the load of the preprocessing module named ashleys-qc-pipeline (see 3.).
Moreover, multiple samples can now be processed during the same execution (https://github.com/friendsofstrandseq/mosaicatcher-pipeline/blob/master/docs/usage.md#classic-behavior) and be part of the same snakemake HTML report.

##### 2.2. External data

Aside from experimental input data, the pipeline also requires a FASTA reference genome. A reference VCF listing Single Nucleotide Variations, for example the 1000 Genomes Project (1000GP; phase 3) (Auton et al. 2015) VCF file can also optionally be used to perform SNPs re-genotyping to detect heterozygous sites from the input data and enable haplotype phasing (Table S1).

##### 2.3. Publishdir

Most of the academic IT infrastructures relies on a combination of different storage technologies, that usually includes persistent hard drives for long-term storage and fast-access flash disks highly coupled to High Performance Computing (HPC) systems where data and softwares sometimes need to be staged first. Additionally, NGS workflows

can produce a high number of intermediate and/or voluminous files, making it difficult to process data efficiently while preserving over the time the relevant outputs computed between space-limited persistent storage and flash disks short-term storage.
To help tackle this problem, MosaiCatcher v2 embeds a “publishdir” new option that allows the user to automatically backup the relevant outputs computed during the execution (read counts table, statistics, GC content, plots, configuration used) in a second predefined location. This allows to preserve both the useful and more heavily used information related to the raw data, as well as the workflow configuration used.

### 3. Strand-seq preprocessing data module (ashleys-qc-pipeline)

As mentioned above, the previous published version of MosaiCatcher took BAM files as input (see above 2. Input data). Strand-seq libraries presenting low coverage (~ 0.03X / cell), different technical issues can arise (list and illustrations here: https://github.com/friendsofstrandseq/MosaiCatcher-pipeline/blob/master/docs/mosaic_count.md). Thus, it is required to identify and flag high-quality cells suitable for processing by using QC plots.

#### 3.1. Ashleys-qc

Following MosaiCatcher publication, a machine learning-based tool named ashleys-qc (Gros et al. 2021) was developed to allow non-expert users to automatically identify low-quality cells based on a series of features including alignment quality and W/C read distribution (see the publication for details).
One main improvement here was to optimise and finetune ashleys-qc snakemake preprocessing pipeline (https://github.com/friendsofstrandseq/ashleys-qc-pipeline) in order to make it both compliant to the snakemake-workflow-catalog standardised usage but also to be usable as a switchable (enabled/disabled) upstream module in MosaiCatcher v2.

Thus, the user can now directly gives as an input to the pipeline the raw Strand-seq FASTQ files to perform a fully automated analysis that goes from the primary processing (QC, mapping, detection of low-quality libraries, GC analysis), secondary analysis (binning, segmentation, haplotype phasing, single-cell SV detection) to tertiary downstream analysis (plots and statistics in an interactive static HTML report).

#### 3.2. Graphic hand-selection of low-quality cells

Additionally to this and as presented on the Figure S1, we developed an alternative mode of execution that allows the user to filter out manually low-quality libraries through a Jupyter Notebook. Practically, after generating sorted and deduplicated BAM files, as well as QC plots, if the “*hand_selection*” parameter is enabled, snakemake will fire an interactive Jupyter Notebook (accessible via a web interface through a SSH tunnel) displaying QC plots next to an interactive sheet (based on ipywidgets and ipysheet modules), allowing the user to unselect low-quality cells (Figure S1). When the resulting output table is produced and the notebook shutdown, the user can launch the same snakemake command without the notebook argument (--edit-notebook) and the pipeline will then continue its processing, based on the content of the new file created. Detailed instructions are provided here.

##### 3.3. Positive control well automatic bypass

To ensure quality control of Strand-seq experiments, it is usually recommended to generate both a positive and a negative control on the 96-well plate. The negative control consists into an empty well, without cells incorporated, while the positive control consists into the introduction of nearly 100 cells, resulting in a much more important coverage for the associated library.
Following ashleys-qc integration into the pipeline, we noticed that the positive control was sometimes labelled as a high-quality library, inducing its processing like a classic single-cell into the SV detection framework. To limit any potential bias related to that, an additional intermediate step was developed to bypass positive control, based on the amount of high-quality usable reads processed by the MosaiCatcher binning method.
This new method, based on z-score outlier detection statistics, takes as input both ashleys-qc predictions for the full plate, as well as the cellwise statistics summary provided by MosaiCatcher binning method, including the binned reads values. To limit the variance, only the cells identified as high-quality libraries by ashleys-qc are then processed, and the z-score applied on the binned reads values. If a z-score value is above 5 (corresponding that the binned reads quantity is deviating from 5 standard deviation from the rest of the distribution), then the well is considered as a positive control.

#### 4. ArbiGent & scNOVA integration

ArbiGent (Porubsky et al. 2022) fork of MosaiCatcher, as well as scNOVA (Jeong et al. 2022) snakemake downstream analysis subworkflow, were cleaned and formatted in order to be compliant with the snakemake-workflow-catalog requirements (conda environment, log files, linting and formatting) as well as with the file structure defined in MosaiCatcher v2.
When enabled, the parameter “*arbigent*” triggers automatic downloading of mappability tracks from Zenodo and loads ArbiGent utility rules as well as the core R program dedicated to SV genotyping and phasing.
In a similar manner, when “scNOVA” parameter is enabled, this one activates automatic downloading of scNOVA Convolutional Neural Networks (CNN) trained models, gene features BED files and triggers scNOVA branch of execution.

#### 5. Continuous Integration/Continuous Development & GitHub actions

As part of the stability increase, MosaiCatcher v2 relies on the GitHub Continuous Integration/Continuous Development (CI/CD) system, named GitHub actions, in order to check pipeline integrity and consistency at each new modification and release. At each new commit, different independent tests are executed, as listed in the Table S2.

In order to reduce potential mistakes and improve code development, new features brought to the pipeline need to be added to the master branch through a pull request, thus requiring to successfully pass these test steps first.

Additionally to these actions, a new version of a docker container including all conda environments and needed dependencies (snakemake –containerize) is automatically built and pushed to a registry when a new tag (git tag) is created.

### 6. Snakemake HTML report

Exploring pipeline generated outputs can be sometimes tedious and difficult looking at the number of different files produced. To tackle this problem, Snakemake can optionally create a static and interactive HTML report that can complement publications and allow better reproducibility and transparency. We leveraged this feature in Mosaicatcher v2 and flagged the outputs listed in Table S3 to be present in the reports (Figure S2). This report allow to embed high-quality figures. Examples are presented in Figure S4 and Figure S5.
The MosaiCatcher report related to data processed in this publication is available as supplemental data.

### 7. Reference genomes

Reference genome is a critical element when it comes to pipeline reproducibility. Indeed, in MosaiCatcher pipeline, reference genome is not only needed to map Strand-seq FASTQ libraries but is used for heterozygous SNP genotyping (without template VCF using bcftools, or with template VCF using freebayes), SNP haplotype phasing (StrandPhaseR) and BAM files haplotagging (whatshap).

To broaden MosaiCatcher pipeline usage, we provide in this new version the possibility to rely on the three main common human genome assemblies: GRCh37/hg19, GRCh38/hg38 (currently the default one) and T2T-CHM13. This can be done using, or not, ashleys preprocessing upstream module (detailed in 3.). To do so, UCSC LiftOver (https://genome.ucsc.edu/cgi-bin/hgLiftOver) and UCSC chain files were used to convert the segmental duplication file from GRCh38 to other assemblies. Remote reference genome Fasta file required is automatically retrieved in MosaiCatcher v2 based on the reference defined by the user (GRCh38 by default) (see Table S4 for sources and registries used).

Finally, as StrandPhaseR relies on the “Software infrastructure for efficient representation of full genomes and their SNPs”, named BSgenome, we adapted BSgenome R reference to the three different assemblies. MosaiCatcher chromosome naming is currently based on UCSC syntax, we are thus automatically referencing the corresponding bioconductor packages referenced in the bioconda registry for assemblies hg19 and hg38. However, no T2T-CHM13 was currently available during the development of MosaiCatcher v2. We thus followed the procedure provided by the BSgenome authors to forge a custom package based on T2T-CHM13 assembly (procedure listed here). The resulting tarball (.tar.gz format) is automatically downloaded and installed in the right conda environment if the user wishes to use T2T assembly (Table S4).

### 8. Data-dependent condition execution snakemake rules

Dealing with pipeline development usually involves thinking about both the normal and expected execution, but also to control specific use-cases or define data-dependent conditions. To handle these behaviours and increase pipeline stability, we defined new "checkpoint" rules based on snakemake feature designed for that purpose (https://snakemake.readthedocs.io/en/stable/snakefiles/rules.html#data-dependent-conditional-execution). While starting from scratch, the user cannot always know which cells would need to be removed, or which chromosomes would potentially present an haploid state. These behaviours previously lead to issues in some cases and needed back and forth running tests and manual correction from the user to avoid them. In MosaiCatcher v2, snakemake "checkpoint" rules allow the user to automatically handle these behaviours by dynamically redefining the Directed Acyclic Graph (DAG) during the pipeline execution (Figure S3).

#### 8.1. High-quality cells filtering

As presented in section 3, the upstream preprocessing module allows the user to identify and flag high-quality cells to be processed in MosaiCatcher v2. To do so, we implemented a "checkpoint" rule that takes as input two files. First, (i) the table produced during the upstream preprocessing module (either by ashleys or by the Jupyter Notebook), representing the list of Stand-Seq libraries and the associated conservative status (to keep or not) of each of these cells. Second, (ii) the output of the mosaic count rule, producing descriptive statistics for each library processed, including a boolean flag indicating if each BAM file (associated to a cell) exhibits sufficient coverage or not. The script used in this rule will thus compute and return as an output, the intersection between the two lists of cells distinctively flagged as high-quality and coverage-sufficient.

Side to this "checkpoint" rule, a python function, used as an input function in snakemake, will read the checkpoint output and return a statement to continue the processing of the direct next steps: count and segmentation at the single-cell level.

#### 8.2. Haploidy detection

One of the strengths of Strand-seq technology is to process and identify chromosomal haploidies at the single-cell level. A natural example of chromosomal haploidy is the single copy of chromosomes X and Y in male human samples. This haploidy status can generate pipeline technical issues, as a key step of MosaiCatcher is to phase heterozygous SNPs genotyped from Strand-seq libraries, implying to distinguish whether these SNPs come from Watson or Crick strands in a diploid chromosomes.

To answer this limitation, we used a snakemake “checkpoint” rule that takes into account ploidy status for all chromosomes. This rule relies on a ploidy estimation python script that processes Watson/Crick read counts in genomic bins of 1Mb to compute a ploidy state estimation for each of these bins across libraries at the sample level. A log-likelihood is computed for each of the ploidy values ranging from 1 to 6, the highest log-likelihood being selected as the ploidy status estimation for the given Mb bin. Median value of ploidy across Mb bins is then computed and used as an input for the checkpoint rule (=1 leads to bypass phasing, >1 leads to phasing).

### 9. SV genotyping through ArbiGent in MosaiCatcher v2

In order to demonstrate ArbiGent capability of genotyping copy-imbalanced SV beyond inversions, we selected previously validated structural variations from the original MosaiCatcher paper (Sanders et al. 2020). Out of the complete list available (https://static-content.springer.com/esm/art%3A10.1038%2Fs41587-019-0366-x/MediaObjects/41587_2019_366_MOESM4_ESM.xlsx), we retained only True Positives (TP) validated through Whole Genome Sequencing (WGS) in the RPE cell lines (RPE-C7, RPE-BM510 and RPE1-WT). As chr10 presents a large-scale 70Mb duplication, we excluded SV present on this chromosome, resulting in a final list composed of 10 copy-imbalanced SV (Table S4). NA20509 LCL sample was kept as a negative control to insure that no RPE-related SV are detected as False Positives. MosaiCatcher was run based on hg38 assembly to match existing validated SV positions. BED file used as input of the pipeline (arbigent_bed_file parameter) can be found here (https://github.com/friendsofstrandseq/mosaicatcher-pipeline/blob/dev/workflow/data/arbigent/paper_backup/manual_segmentation_validation_paper.bed).

### 10. Multistep read count normalisation

Adjusting for library size and library composition follows a median of ratios approach (Anders and Huber 2010). Briefy, a pseudo-reference sample is calculated from the geometric mean of read counts per bin, across cells, including only bins where all libraries have non-zero counts. Then, scaling factors are calculated as the median of the ratios of bin read counts relative to the pseudo-reference sample.
The relationship between GC content and per-bin read counts is derived with the locally weighted scatterplot smoothing (lowess) algorithm. In order to speed up calculations, the lowess curve is fitted only to a subset of the read counts, subsampled within GC content deciles. Finally, from the fitted curve, per-bin correction factors are calculated depending on their GC content.
Similarly to RNA-seq and other count-based data, Strand-seq reads follow a negative binomial distribution. Therefore, it is possible to find a variance stabilising transform for the data for visualisation purposes, such as the Anscombe transform (ANSCOMBE 1948).

Variance stabilising transformation requires the estimation of dispersion, which done by minimising the estimated residual variance (Harrison 2017) .

### Supplementary data

### 1. Supplemental tables

| Name | Description | Questions |
|---|---|---|
| `Formatting` | Files formatting (snakefmt) | Did the formatter identify some aesthetic potential improvements to the code? |
| `Linting` | Files linting (snakemake –lint) | Did the linter identify functional improvements, missing declarations or code ambiguities? |
| `Linting_ashleys` | Files linting (snakemake –lint) with ***ashleys_pipeline*** enabled | |
| `Linting_arbigent` | Files linting (snakemake –lint) with ***arbigent*** enabled | |
| `Testing` | Pipeline execution | Did the pipeline run without errors using the smoke dataset provided under different conditions?<br>Were there any errors during the creation of the HTML report? |
| `Testing_norm_disabled` | Pipeline execution with HGSVC-based reads count **normalisation** disabled | |
| `Testing_ashleys` | Pipeline execution with ***ashleys_pipeline*** enabled | |
| `Testing_ashleys_HGSVC_norm_disabled` | Pipeline execution with HGSVC-based reads count **normalisation** enabled with ***ashleys_pipeline*** enabled | |
| `Testing_ashleys_mulsitep_normalisation_correction_enabled` | Pipeline execution with ***ashleys_pipeline*** enabled and multistep normalisation module enabled | |
| `Testing_ashleys_jup_nb` | Pipeline execution with ***ashleys_pipeline*** enabled and use jupyter notebook skeleton instead of ashleys-qc to check dependencies | |
| `Testing_ashleys_norm_enabled_hg19` | Pipeline execution with ***ashleys_pipeline*** enabled using full hg19 fasta file needed to be downloaded and indexed instead of chr17 fasta present in the git repository | |
| `Testing_ashleys_norm_enabled_hg38` | Pipeline execution with ***ashleys_pipeline*** enabled using full hg38 fasta file needed to be downloaded and indexed instead of chr17 fasta present in the git repository | |
| `Testing_ashleys_norm_enabled_T2T` | Pipeline execution with ***ashleys_pipeline*** enabled using full T2T fasta file and BSgenome package needed to be downloaded and indexed instead of chr17 fasta present in the git repository | |
| `Testing_arbigent` | Pipeline execution with ***arbigent*** enabled, including automatic downloading of mappability track from Zenodo | |

| | |
|---|---|
| `Testing_publishdir` | Pipeline execution with ***publishdir*** option enabled |

*Table S1 - GitHub actions defined in the MosaiCatcher v2 git repository*

| **Output** | **Format type** | **Comment** | **Ashleys-qc / MosaiCatcher** |
|---|---|---|---|
| FastQC | HTML | Quality Control of Strand-seq FASTQ libraries | **Ashleys-qc** |
| Mosaic QC plot | PDF | Reads density across bins: allow to visualise both global statistics at the sample level but also individual Strand-seq karyotypes at the cell level | **Ashleys-qc/ MosaiCatcher** |
| Mosaic QC plot - after multistep normalisation | PDF | | |
| GC correction plot | PNG | Scatter plots to represent data GC composition before (left) and after (right) GC correction during the multistep normalisation. The lowess smoothing line helps to identify correlation between GC composition and read counts in a representative subset of the data. | **Ashleys-qc** |
| VST histogram | PNG | Histogram representation of the read count distribution across bins before (left) and after applying the Variance Stabilising Transformation method, part of the multistep normalisation. | **Ashleys-qc** |
| Plate plot | PNG | Plot representation of ashleys-qc probabilities/binary-predictions. Can allow the user to detect more easily an experimental issue. | **Ashleys-qc** |
| SV calls | PDF | Representation at the chromosome level of the SV landscape for each individual library that has been able to be processed. | **MosaiCatcher** |
| SV clustering | PDF | Clustered heatmap representation of the SV events that allow the user to have a complete and global representation at the sample level using Ward Hierarchical Agglomerative clustering. Two versions of the heatmap are provided: one scaled according to the chromosome size and one unscaled, based on the event presence/absence. | **MosaiCatcher** |
| SV consistency | PDF | Barplots representing SV events (rows) according to their frequency/position across cells and their class (del, dup, inv, ...) | **MosaiCatcher** |
| Run summary | TXT | Summary of the run including: configuration used, cells that | **MosaiCatcher** |

| | | |
|---|---|---|
| | | were processed based on both ashleys/hand-selection and coverage compliancy, chromosomes that were phased based on the predicted ploidy status. |

*Table S2 - Outputs referenced in the MosaiCatcher v2 HTML report*

| **Collection** | **Sample name** | **Processed / Sequenced libraries** | **Positive control** | | | | **Bypassed haploid chromosome(s) phasing** |
|---|---|---|---|---|---|---|---|
| | | | **Library** | **ashleys-qc probability** | **Nb of reads used in mosaicatcher** | **z-score** | |
| RPE cell lines | RPE-BM510 | 82/96 | BM510x3PE20451 | 0.2462 | / | / | chr13 |
| | RPE1-WT | 91/96 | RPE1WTPE20488 | 0.6348 | 16,942,496 | 9.44 | / |
| | RPE-C7 | 86/96 | C7x02PE20315 | 0.7753 | 48,366,179 | 9.21 | chr13 |
| Healthy HGSVC sample | NA20509 LCL | 53/96 | GM20509Bx01PE20515 | 0.6232 | 1,979,346 | 5.35 | chrX,chrY (male sample) |

*Table S3 - Samples previously processed into a unique run*

window size: 200kb ; assembly: T2T-CHM13

| ID | Length (kb) | SV type ground truth | NA20509 LCL | RPE-C7 | RPE-BM510 | RPE1-WT | % genotype concordance (compared to validated genotype) |
|---|---|---|---|---|---|---|---|
| chr3-60900000-62300000 | 1400 | dup | 0\|0 | 0\|0 | **1020** | **2010** | **100%** |
| chr3-102800000-103600000 | 800 | del | 0\|0 | **1000** | 0\|0 | 0\|0 | **100%** |
| chr9-21900000-22400000 | 500 | del | 0\|0 | 0\|0 | **1000** | **0010** | **100%** |
| chr12-25100000-25800000 | 700 | dup | 0\|0 | 0\|0 | **2010*** | 0\|0 | **100%** |
| chr15-93300000-101991189 | 8691.189 | idup/inv | 0\|0 | **2110*** | **0020*** | 0\|0 | **100%** |
| chr16-77600000-78100000 | 500 | dup | 0\|0 | 0\|0 | **1020** | **3010*** | **100%** |
| chr17-63100000-63500000 | 400 | del | 0\|0 | 0\|0 | **1000** | **0010** | **100%** |
| chr20-3900000-4100000 | 200 | del | 0\|0 | 0\|0 | **1000** | **1000** | **100%** |
| chr22-37900000-39700000 | 1800 | idup | 0\|0 | 0\|0 | **2110*** | 0\|0 | **100%** |
| chr22-39700000-40400000 | 700 | dup | 0\|0 | 0\|0 | **2110*** | 0\|0 | **100%** |

*Table S4 - Genotyping of 10 copy-imbalanced validated SV through ArbiGent mode in MosaiCatcher v2*

We demonstrate ArbiGent capability of genotyping copy-imbalanced SV using a list of 10 events previously validated through Whole Genome Sequencing (WGS). 0|0 corresponds to absence of the SV. Haplotype code of consensus call [ 0101 : H1 ref orientation copies; H1 inverted orientation copies; H2 ref orientation copies; H2 inverted orientation copies]. Haplotype codes followed by an asterisk (*) indicate that the same SV was detected but with a slight modification compared to the original result (3010 instead of 2010 for chr16-77600000-78100000 in RPE1-WT for instance).

| Assembly | Name | URL | Default |
| --- | --- | --- | --- |
| GRCh38/hg38 | FASTA | https://hgdownload.soe.ucsc.edu/goldenPath/hg38/bigZips/analysisSet/hg38.analysisSet.fa.gz | X |
| | R BSGenome | https://bioconductor.org/packages/release/data/annotation/html/BSgenome.Hsapiens.UCSC.hg38.html | |
| GRCh37/hg19 | FASTA | https://hgdownload.soe.ucsc.edu/goldenPath/hg19/bigZips/analysisSet/hg19.p13.plusMT.no_alt_analysis_set.fa.gz | |
| | R BSGenome | https://bioconductor.org/packages/release/data/annotation/html/BSgenome.Hsapiens.UCSC.hg19.html | |
| T2T-CHM13 | FASTA | https://s3-us-west-2.amazonaws.com/human-pangenomics/T2T/CHM13/assemblies/analysis_set/chm13v2.0.fa.gz | |
| | R BSGenome** | https://zenodo.org/record/7697400/files/BSgenome.T2T.CHM13.V2_1.0.0.tar.gz?download=1 | |

*Table S5 - Reference genome files external data*

### 2. Supplemental figures

A

B

C

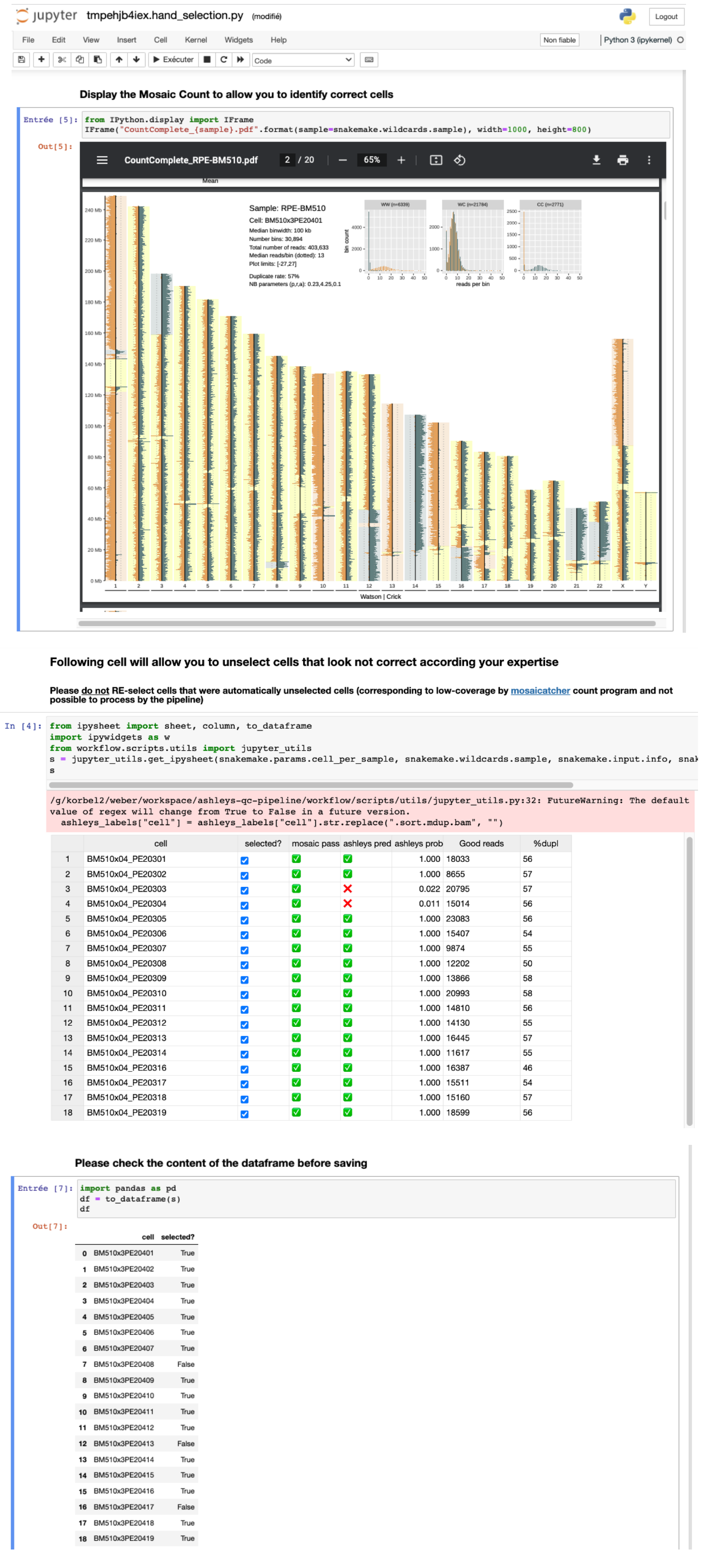

*Figure S1 - Manual filter-out of low-quality cell using a web interface*

Screenshot of the jupyter notebook interface. As explained in the documentation here, snakemake allows you to fire and edit a jupyter notebook using the right parameters. Thus, (A) the user here can visualise the Strand-Seq karyotypes representing the density and the Watson/Crick reads distribution across the chromosomes in order to (B) unselect low-quality libraries based on his expertise. The interactive table is then transcribed into a pandas dataframe (C) and exported as a TSV table, required for the following part of the analysis.

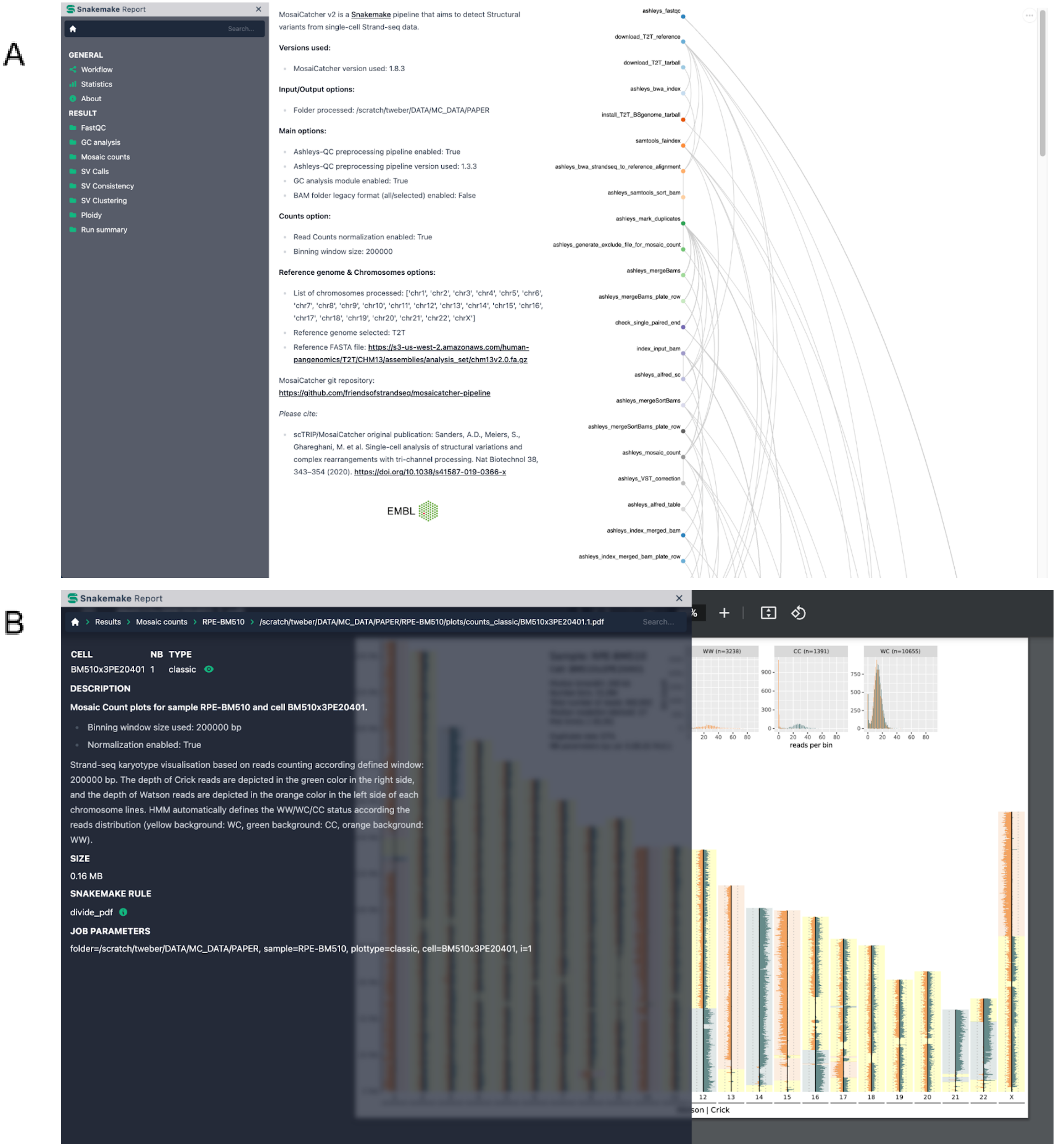

*Figure S2 - MosaiCatcher v2 report screenshots*

(A) Home page of MosaiCatcher v2 HTML report produced. Home page contains both config parameters used when executing the run but also the DAG. (B) Example of visualisation for the category "Mosaic counts", representing a Strand-Seq karyotype at the cell level. Users have access both to the visualisation but also to dynamic description that can be adapted using ReStructuredText and snakemake wildcards.

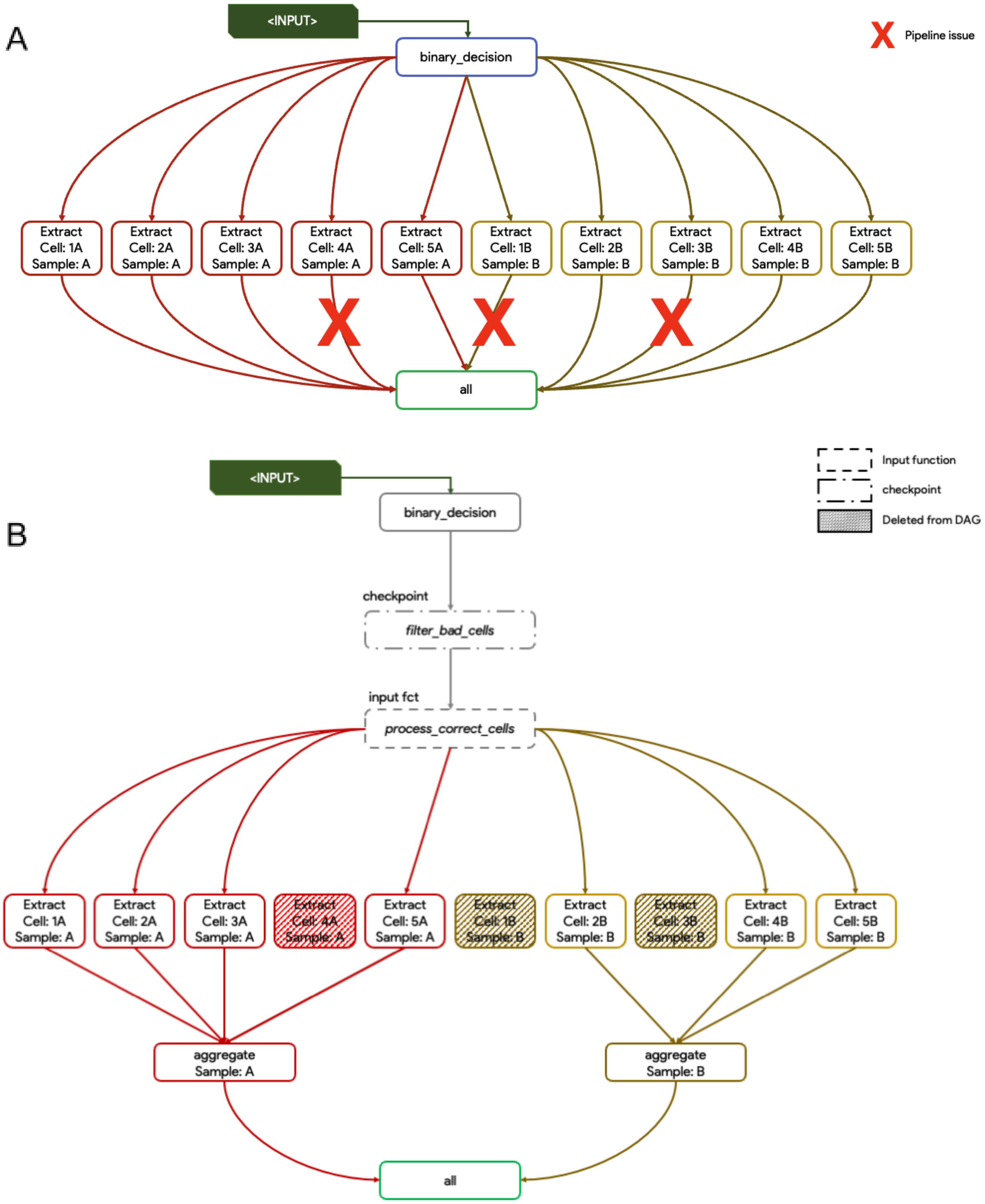

*Figure S3 - schematic representation of snakemake checkpoint rule*

(A) Based on a set of input, here 2 samples (Sample A & Sample B), each containing 5 cells are processed through a snakemake pipeline without knowledge about the cells quality, leading to a pipeline issue. (B) Using a snakemake checkpoint rule, low-quality cells can be automatically removed during the pipeline execution by modifying the DAG dynamically.

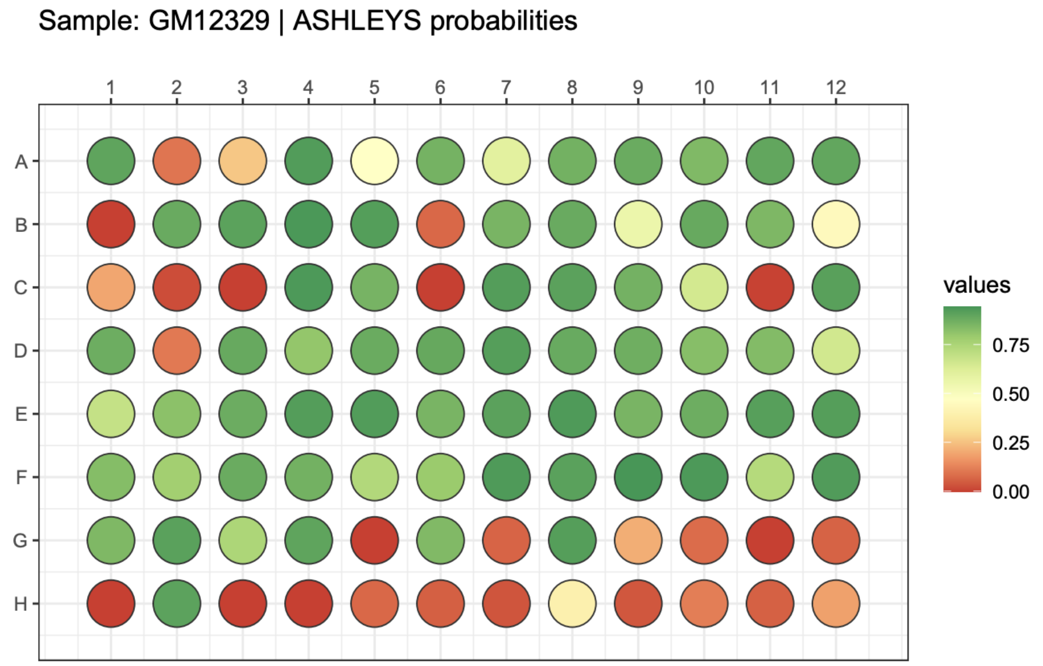

*Figure S4 - Ashleys probabilities in a 96-well plate visualisation*

Ashleys-qc probabilities and binary predictions can be visualised through a plot representation that allows to identify more easily potential contamination or experimental issues.

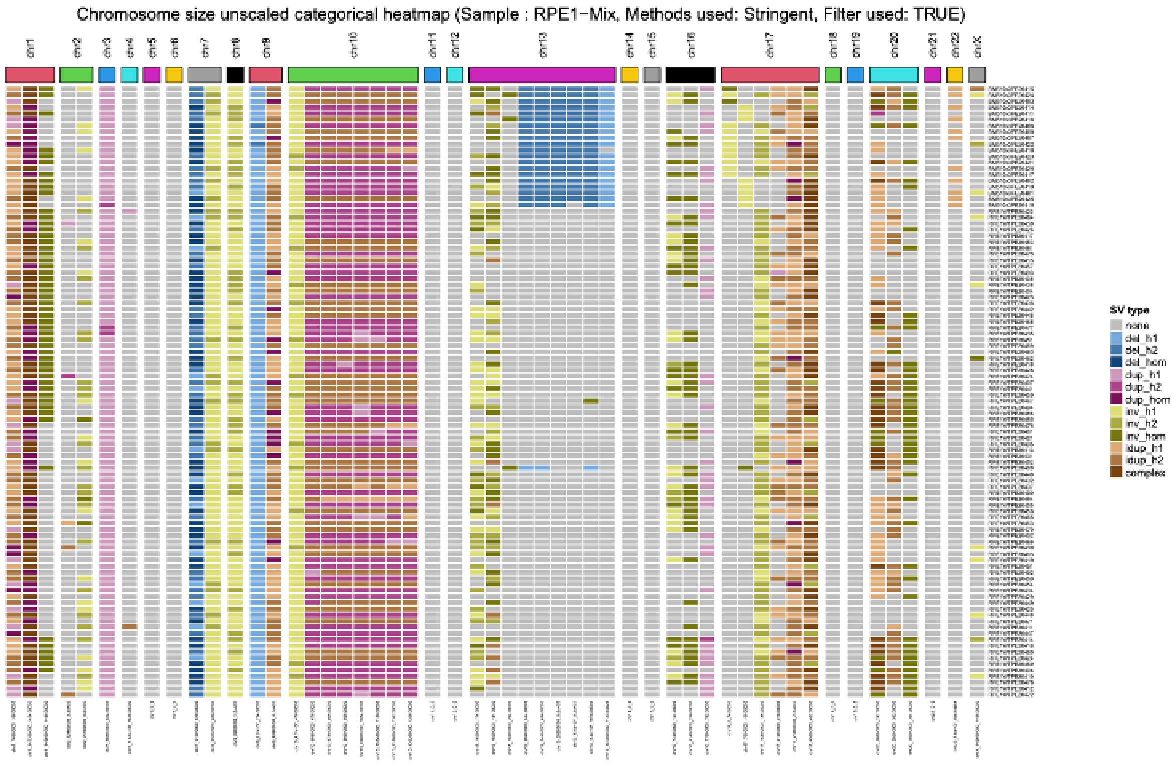

*Figure S5 - SV distribution heatmap visualisation update*

An example of updated and improved visualisation in MosaiCatcher v2. This heatmap represents the phased SVs called into the plate according to their types (deletion (del), duplication (dup), inversion (inv), inverted duplication (idup) and complex event) along the chromosomes (in column) and the presence/absence in the different libraries sequenced (in rows).

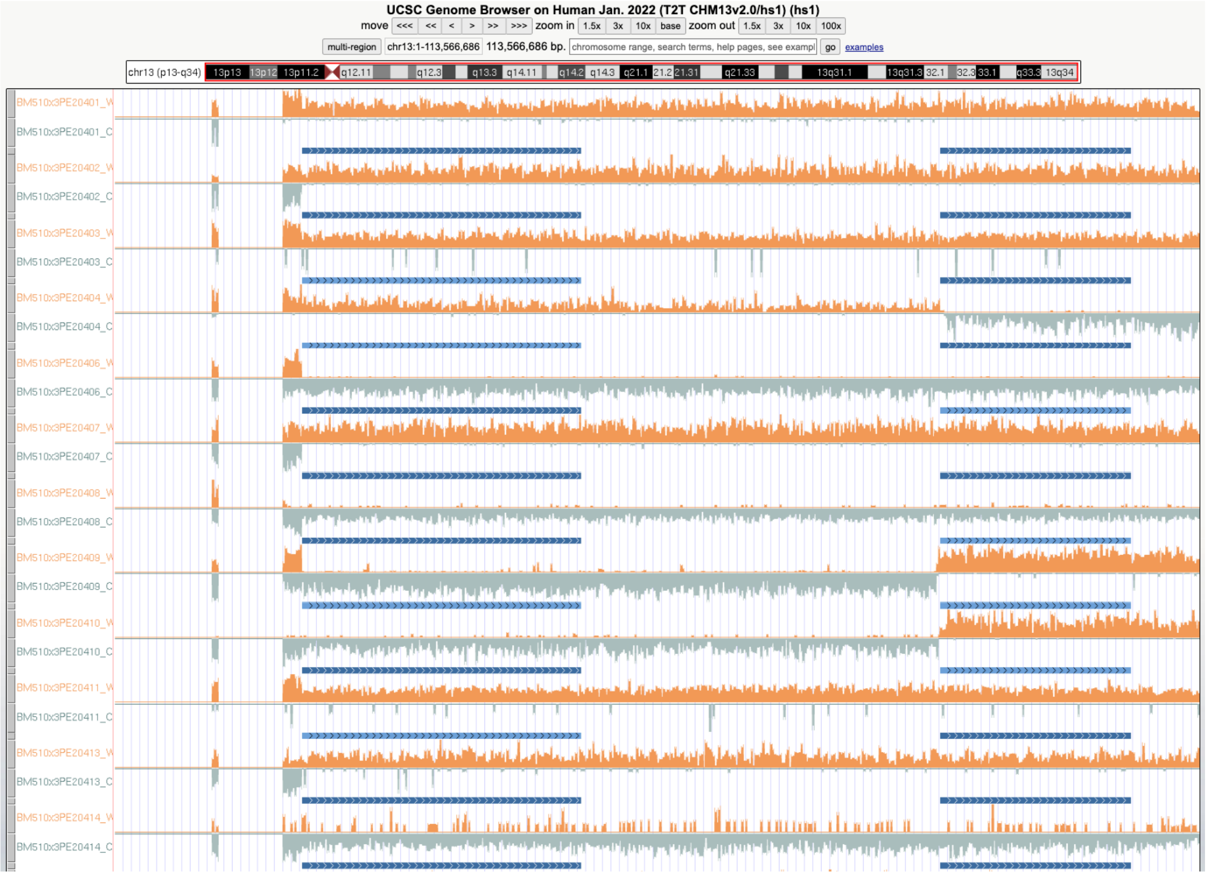

*Figure S6 - UCSC genome browser visualisation of data produced by MosaiCatcher v2*

Example of Strand-Seq data visualisation in the UCSC browser, using file generated by MosaiCatcher v2. Cells are displayed in the alphabetical order. 3 tracks are available for each cell: the Watson read counts, the Crick read counts and the SV calls detected at the cell level. Here is an example of a clonal phased deletion event detected on chr13 of the RPE-BM510 sample. By hovering the mouse over read counts or SV calls, additional information is displayed.
